# Structural basis of microtubule destabilization by GTP hydrolysis

**DOI:** 10.64898/2026.06.19.733065

**Authors:** Juan Estévez-Gallego, Pavel Filipcik, Hugo Muñoz-Hernández, Senik Matinyan, Michal Wieczorek, Michel O. Steinmetz

## Abstract

Microtubules are cytoskeletal filaments that dynamically grow and shrink to support vital biological functions. Their behavior is regulated by GTP hydrolysis in tubulin incorporated into the lattice, but the underlying chemical transitions and their impact on microtubule stability remain unclear. Using cryo-electron microscopy, we resolved microtubule structures at 1.9–2.2 Å in four different nucleotide states, revealing the roles of amino acids, ions, and hundreds of water molecules in the hydrolysis process. We found that compaction of the GTP-bound lattice into a more stable pre-hydrolysis state activates a catalytic water molecule for γ-phosphate cleavage. Hydrolysis intermediates are stabilized by strengthened lattice contacts, but in the final GDP state, release of reaction products disrupts these contacts and destabilizes the lattice. Our results explain how chemical changes in tubulin’s nucleotide site drive lattice transitions and offer an atomistic framework for understanding microtubule dynamic instability.

## Main Text

Microtubule dynamic instability is fundamental to eukaryotic cell biology (*1, 2*), which is driven by GTP hydrolysis within the microtubule lattice (*3–5*). Microtubules are non-covalent polymers of αβ-tubulin heterodimers (hereafter tubulin) that assemble by establishing longitudinal and lateral interactions within the microtubule lattice (*5, 6*). The stochastic switching between microtubule growth and shrinkage, termed dynamic instability, is a defining feature of microtubule behavior essential for critical eukaryotic cellular functions such as chromosome segregation, cell migration, intracellular transport, and mechanochemical signaling (*7–9*). This behavior is governed by the hydrolysis of GTP to GDP at growing microtubule ends, which triggers disassembly through the loss of a stabilizing, GTP-tubulin containing region known as the GTP cap (*1, 3, 10*). While GTP hydrolysis occurs slowly in free tubulin and in tubulin oligomers (*11, 12*), it is accelerated nearly a thousand-fold upon incorporation into the microtubule lattice (*13, 14*). Within the lattice, the α-subunit of one tubulin dimer contributes catalytic residues that, together with the exchangeable nucleotide site (E-site) of the β-monomer from an adjacent dimer, completing the catalytic site at the longitudinal interdimer interface. Consequently, it has been proposed that the microtubule lattice confers a GAP-like activity to accelerate tubulin GTP hydrolysis (*15–17*).

In current models, GTP hydrolysis generates lattice strain that destabilizes microtubules, ultimately triggering catastrophe - that is, the switch from microtubule growth to shrinkage (*18–21*). However, the precise mechanism underlying this process, as well as the role of water molecules in coupling the hydrolysis reaction to lattice destabilization, remains elusive. This knowledge gap stems largely from the resolution limits (∼3-4 Å) of existing native, undecorated cryo-electron microscopy (cryo-EM) microtubule structures (*20–22*). At these resolutions, it is not possible to accurately model solvent molecules or the exact orientations of amino acid side chains. As a result, the roles of water molecules and the Mg²⁺ ion at the catalytic site, as well as how water-and side chain-mediated networks influence longitudinal and lateral tubulin–tubulin interactions within the lattice, remain unclear. Determining these structural features is essential for a comprehensive understanding of how chemical transitions during GTP hydrolysis dictate microtubule dynamic instability.

### Sub-2.2 Å resolution structures of microtubules in four nucleotide states

To address this knowledge gap, we used cryo-EM to determine six structures of 13- and 14-protofilament (pf), 3-start bovine brain microtubules at resolutions between 1.9 and 2.2 Å. Our structures capture four distinct states of the microtubule lattice-associated GTPase reaction using well-established nucleotides or their analogs (*23–26*) (Table S1-S3): GMPCPP (13-pf, 2.2 Å; 14-pf, 1.9 Å), GDP·BeF₃^-^ (13-pf, 2.0 Å), GDP·AlF₃ (13-pf, 2.0 Å), and GDP (13-pf, 2.2 Å; 14-pf, 2.0 Å). Here, BeF₃^-^ and AlF₃ serve as inorganic γ-phosphate (P_i_) analogs that represent the tetrahedral ground state and a planar hydrolysis intermediate, respectively (*24–26*). The quality of the density maps enabled accurate modeling of side chains, ligands, and ions, and, remarkably, hundreds of ordered water molecules distributed throughout the microtubule lattice (Fig. 1, Table S4, Figs S1 and S2). Water molecules were modeled only when supported by a clear density and a chemically plausible local environment; their high average Q-scores of >0.84 across all structures confirm the reliability of their placement (*27*).

**Figure 1.**
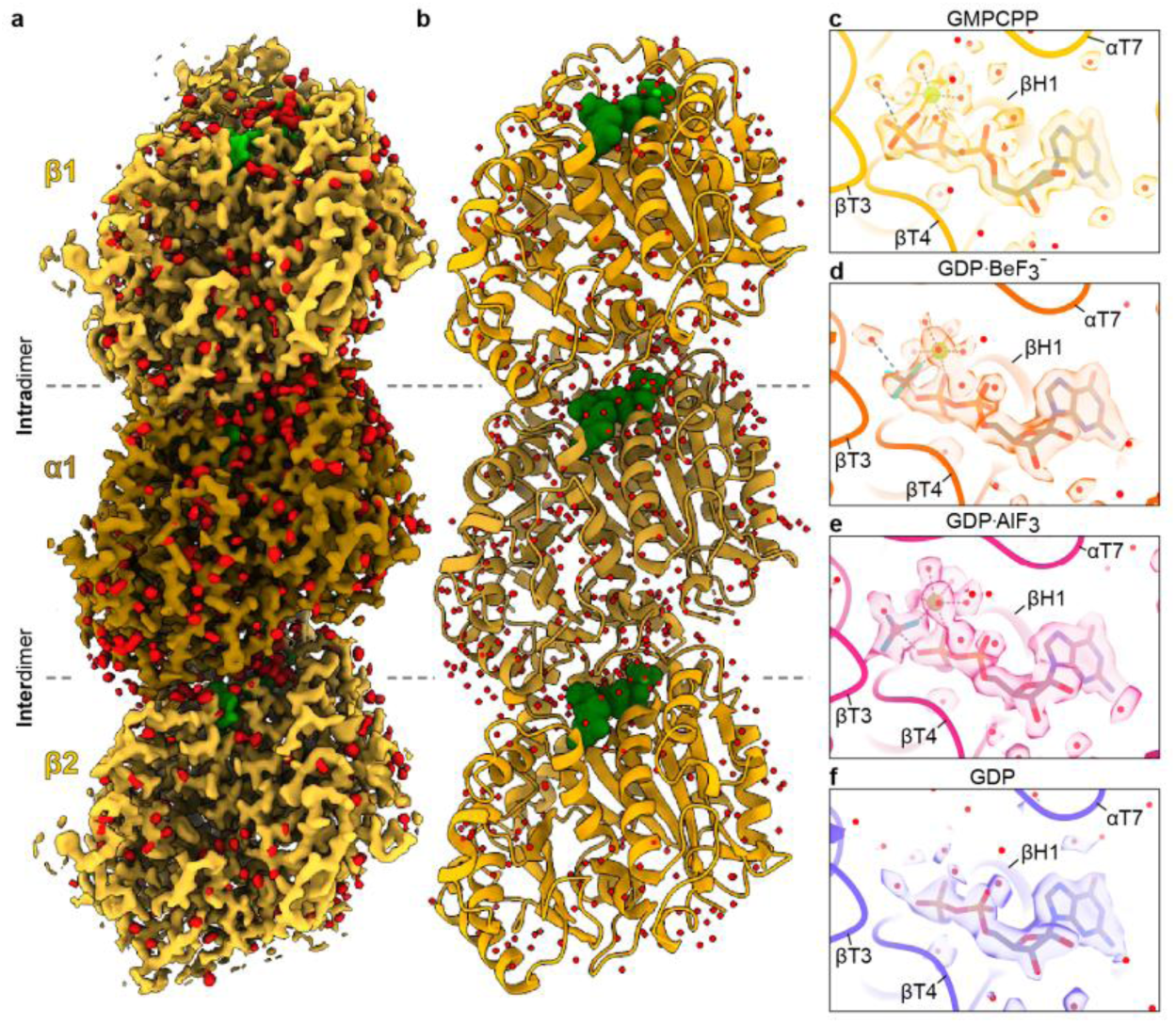
Sub-2.2 Å resolution microtubule structures. (**a**) Luminal view of the cryo-EM density map of a GMPCPP-microtubule protofilament, with protein, ligands and waters shown in yellow, green, and red, respectively. Densities are contoured at two times root mean square deviations (2 RMSD; protein and ligand) and 1 RMSD (water). (**b**) Cartoon representation of the model of the 14-pf GMPCPP-microtubule protofilament with waters and Mg^2+^ ions shown as red and lime spheres, respectively. The nucleotides are shown in green spheres representation. (**c-f**) Close-up views of catalytic sites with models and density maps of nucleotides (stick representation), water molecules (red spheres), and Mg^2+^ ions (green spheres) shown at 2 RMSD. Panels illustrate: (**c**) 14-pf GMPCPP-(yellow, 1.9 Å resolution), (**d**) 13-pf BeF_3_^-^-(orange, 2.0 Å resolution), (**e**) 13-pf AlF_3_-(magenta, 2.0 Å resolution), and (**f**) 14-pf GDP-microtubules (blue, 2.0 Å resolution).

Our models reveal distinct conformational transitions across the microtubule lattice during GTP hydrolysis (Table S5, Movie S1). As illustrated for the 13-pf reconstructions in Fig. 2, GMPCPP-microtubules display an expanded lattice with a right-handed pf twist (dimer rise and pf twist of 84.9 Å and 0.17°, respectively). In contrast, GDP·BeF₃^-^- and GDP·AlF₃-microtubules exhibit a compacted lattice with a left-handed pf twist (dimer rise and pf twist of 82.3 Å and −0.07°, and 82.3 Å and −0.08°, respectively). Finally, GDP-microtubules maintain a compacted lattice but display nearly straight pfs (dimer rise and pf twist of 82.4 Å and 0.04°, respectively). While transitions from expanded to compacted states and from right-handed to left-handed twists have been described previously in undecorated microtubules (*20–22*), our high-resolution data provide the structural foundation to correlate these global lattice rearrangements with the specific chemical state of the nucleotide.

**Figure 2.**
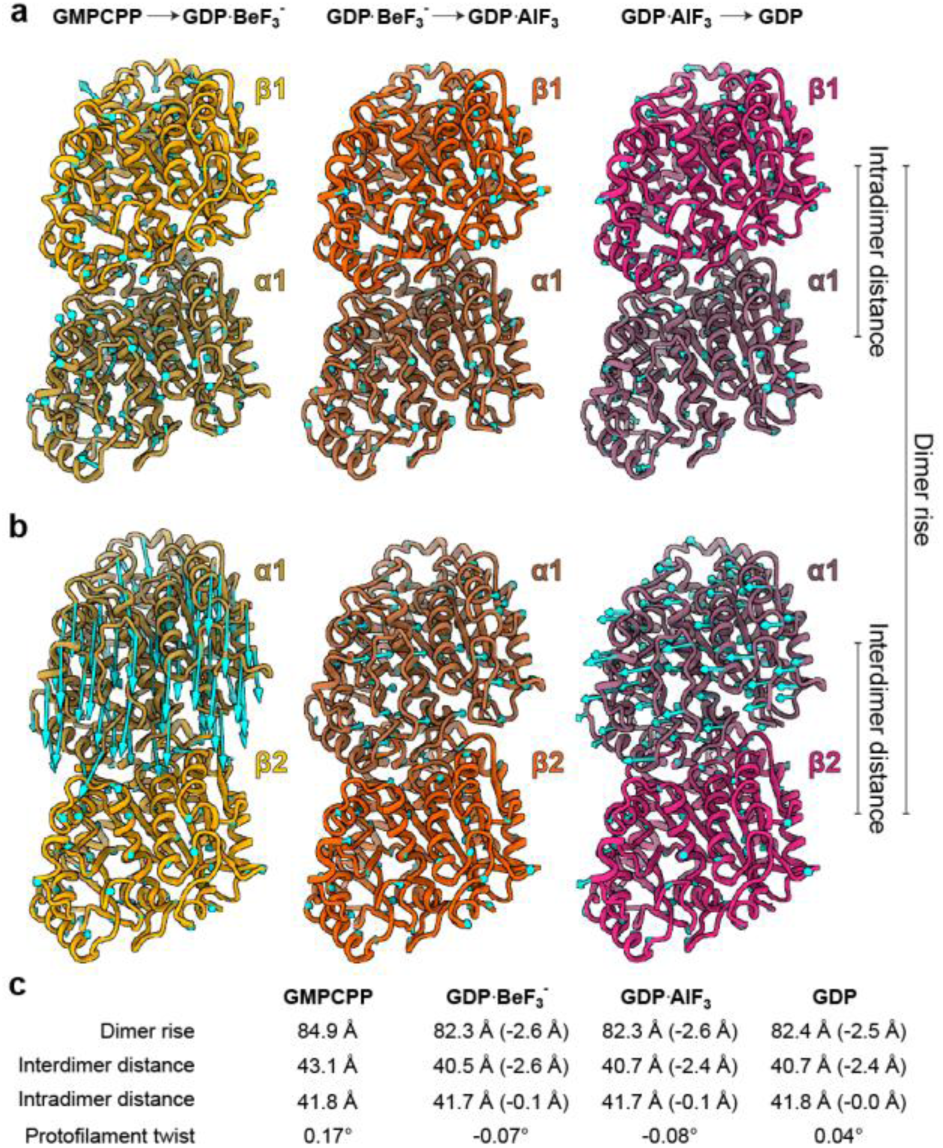
Microtubule lattice conformational transitions during GTP hydrolysis. Cartoon representation of GMPCPP-(yellow), GDP·BeF₃⁻ (orange), and GDP·AlF₃ (magenta) tubulin states from their corresponding trimer models. Panels (**a**) and (**b**) show the intradimer and interdimer regions, respectively. Cyan arrows represent conformational displacement vectors derived from structural comparisons between states, calculated after alignment of dimers using the N-terminal domain of α-tubulin (upper panel) or β-tubulin (middle panel) as reference. Arrow lengths are depicted 5-fold larger and every 5th Cα displacement is shown for clarity. Panel (**c**) presents a table summarizing lattice parameters for each 13-pf microtubule reconstruction. An extended version of the table, including 14-pf reconstructions is included as Table S5.

### Molecular mechanism of GTP hydrolysis

The resolutions of our 13- and 14-pf microtubule reconstructions allowed us to deduce a molecular mechanism for GTP hydrolysis within the tubulin catalytic site. From here onwards, secondary structure elements in mammalian tubulin are defined according to established conventions (*5, 28*) (helices, H; strands, S; loops, T or two secondary structure elements connected by a hyphen) and with residues and secondary structure elements assigned to their α- or β-tubulin monomer. Movie S2 shows the transitions between the four nucleotide states in the active site.

The GMPCPP-, GDP·BeF₃^-^-, and GDP·AlF₃-microtubule structures all contain a water molecule that is well-positioned to serve as a catalytic nucleophile (Fig. 3a–c, Fig. S3aii-cii). This water is stably coordinated through hydrogen bonds with the side chains of residues αGlu254 and αAsp251 in loop αT7, and the main chain amide of βGly100 in loop βT3. A Mg²⁺ ion is also present in all nucleotide analog structures. It adopts an octahedral coordination with four water molecules and two oxygen atoms from the nucleotide β-phosphate, and the γ-phosphate or its analogs (Fig. 3a–c, Fig. S3ai-ci). These coordinating water molecules form variable hydrogen bonds with αAsp251, αGlu254, βGln11, βAsp69, βGlu71, βThr74, and βThr145 across the different structures. In addition, the oxygen and fluorine atoms of the γ-phosphate and its analogs, respectively, engage in direct-or water-mediated hydrogen bonds with αAsn249, αAsp251, αGlu254, βAla99, βGly100, βAsn101, βGly144, and βThr145 (Fig. S3aii-cii). In the GDP-microtubule structure, the Mg²⁺ ion and the γ-phosphate are absent, and the ordered water molecules are reorganized (Fig. 3d, Fig. S3d).

**Figure 3.**
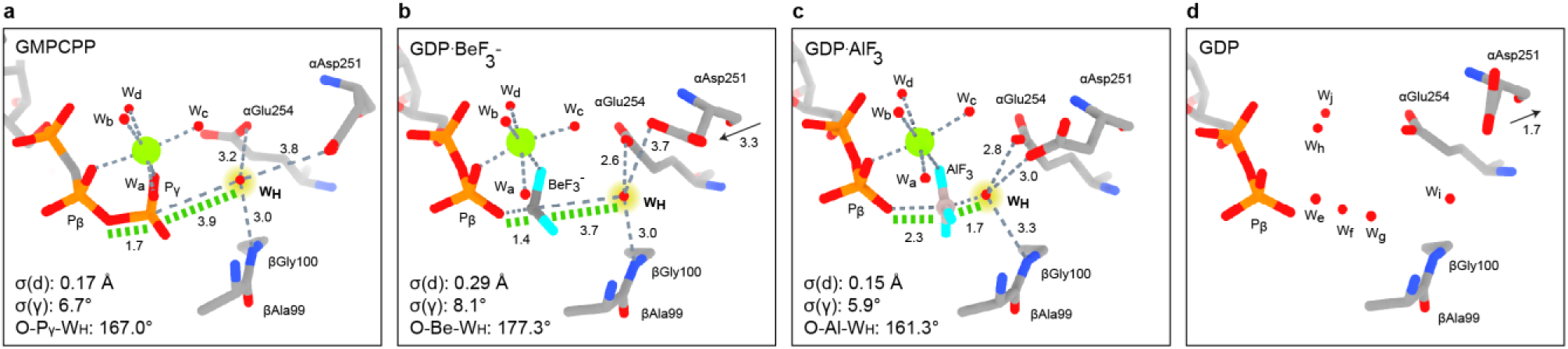
Snapshots of the GTP hydrolysis mechanism. Close-up views of the catalytic site in GMPCPP (**a**), GDP·BeF₃⁻ (**b**), GDP·AlF₃ (**c**), and GDP (**d**) microtubules. Water molecules (w) are depicted as small red spheres, with catalytic water emphasized with a yellow rim (w_H_). The Mg²⁺ ion is shown as larger green spheres. Distances are illustrated as dashed lines and reported in Å. The angle of the catalytic water relative to both the γ-phosphate and the β–γ bridging oxygen is given and represented with a green dashed line. The standard deviation of the bond lengths of (σ_d_) and angles (σ_γ_) of the waters forming the Mg²⁺ coordination sphere (w_a_-w_d_) are given.

We propose that the interplay of these elements drives the cleavage of the nucleotide’s β–γ phosphate ester bond. To facilitate GTP hydrolysis, the catalytic water molecule must approach the γ-phosphate to a near-covalent distance (<2 Å) and adopt an angle close to 180° relative to the ester bond axis for optimal nucleophilic attack (*29*). In the expanded GMPCPP-microtubule lattice, this optimal geometry is not yet achieved: the catalytic water remains at a relatively long distance (3.9 Å) from the γ-phosphate and is oriented at a suboptimal angle (167.0°; Fig. 3a).

In contrast, compaction of the lattice prior to GTP hydrolysis yields a pre-hydrolysis state represented by the GDP·BeF_3_-microtubule structure, in which loop αT7 shifts toward the nucleotide. This movement brings αGlu254 closer to the catalytic water (from 3.2 to 2.6 Å) and substantially shifts αAsp251 by 3.3 Å toward the catalytic site, accompanied by a change of its side chain conformation (Fig. 3b). Consequently, the catalytic water adopts a more favorable geometry for nucleophilic attack, forming an angle of 177.3° with the β–γ phosphate ester bond. In parallel, the Mg²⁺ coordination sphere becomes distorted, displaying increased variability in water coordination distances and angles (Fig. 3b, Fig. S3bi).

Once the catalytic water is properly positioned, the hydrolysis reaction proceeds via a GDP·P_i_ transition state, represented by the GDP·AlF₃-microtubule post-hydrolysis structure. Three factors promote the formation of this state. First, the catalytic αGlu254 residue (*16, 30*) initiates the deprotonation of the catalytic water molecule for nucleophilic attack (*14*). Second, the side chain of αAsp251 undergoes a conformational change, positioning the catalytic water at a covalent distance (from 3.7 to 1.7 Å) from the γ-phosphate (Fig. 3c). This observation provides a structural basis for the role of αAsp251 in GTP hydrolysis, whose function has previously been inferred based on mutagenesis (*31*). Lastly, the Mg²⁺-water coordination sphere relaxes back to a regular octahedral geometry (*32*) (Fig. 3c, Fig. S3ci). This relaxation facilitates a partial dissociation of the γ-phosphate from the β-phosphate, as evidenced by an expansion of its bond length from 1.4 to 2.3 Å (Fig. 3c, Fig. S3cii). Collectively, these factors promote the formation of a pentavalent phosphorane intermediate, in which the catalytic water forms a covalent bond with the γ-phosphate to facilitate cleavage of the β–γ phosphate ester bond (*33*).

Following hydrolysis, the transition to the final GDP-microtubule state involves the release of P_i_ and the Mg²⁺ ion. This triggers a dramatic reorganization of the catalytic site, in which the side chain of αAsp251 is repositioned, and the newly cleared space within the catalytic site is subsequently filled by ordered water molecules (Fig. 3d, Fig. S3d).

### Consequences of GTP hydrolysis for lattice stability

Our structures reveal that GTP hydrolysis is accompanied by extensive remodeling of tubulin–tubulin interactions at both the longitudinal (α_1_-β_1_ intradimer and β_1_-α_2_ interdimer) and lateral (α–α’ and β–β’) interfaces (Fig. 4a; subscripts 1 and 2 denote dimer identity; the prime indicates a tubulin monomer from a neighboring pf). We quantified these interface dynamics by defining contacts as residue pairs linked by direct or water-mediated interactions (Fig. 4b-e, Figs S4-S8). As illustrated in Fig. 4b, from the initial expanded GMPCPP-to the final compacted GDP-microtubule states, longitudinal interfaces are extensively reorganized (Fig. S8ab) but show only a modest net increase in total contacts. In contrast, we observed a significant loss of lateral contacts, particularly those mediated by the αM-loop (Fig. S8c). These findings support a model where lattice destabilization is primarily driven by the weakening of lateral, rather than longitudinal interactions (*19*).

**Figure 4.**
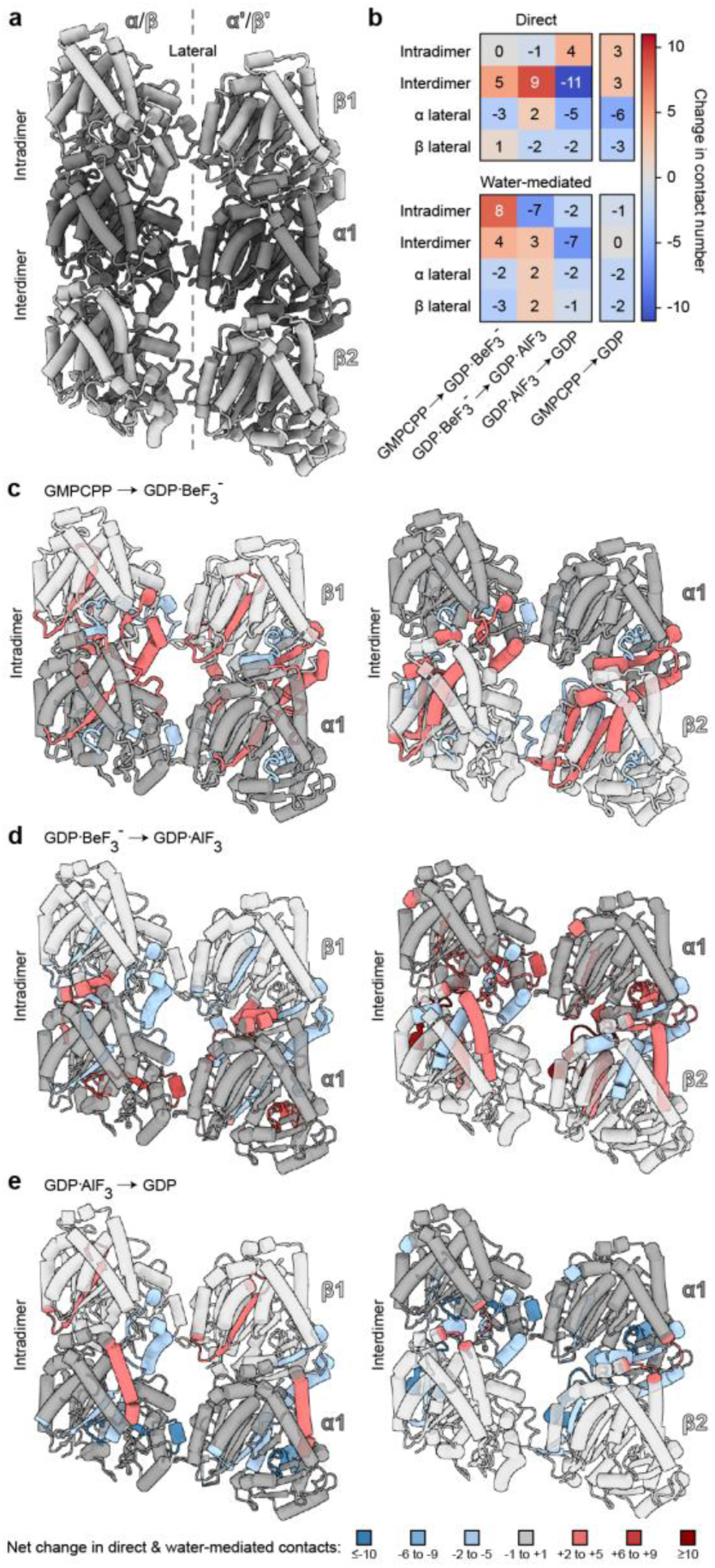
Microtubule lattice rearrangements during GTP hydrolysis. (**a**). Cartoon representation of two tubulin trimers indicating the interfaces defined for the contact analysis. α-tubulin is shown in light gray and β-tubulin is shown in dark gray. The lateral interface is indicated by a dashed vertical line. (**b**). Heatmaps depicting the changes in direct (upper panel) and water-mediated (lower panel) number of contacts at the different interfaces between the hydrolysis reaction stages. The last column shows the overall changes between the initial GTP-expanded (GMPCPP) and final (GDP) stages. The color key indicates the correspondence between color and changes in contact numbers. (**c–e**). Representation of the changes in number of contacts for each secondary structural element between the different stages of the hydrolysis reaction. Cartoon representations of two tubulin dimers, centered at the intradimer interface (left) and four neighboring tubulin monomers, centered at the interdimer interface (right). The color key is shown at the bottom of the figure, indicating the correspondence between color and net changes in contact numbers. α-Tubulin is shown in light gray; β-tubulin is shown in dark gray. Regions that are not colored with net changes in contacts are transparent for clarity.

Examination of the tubulin-tubulin interfaces through each stage of GTP hydrolysis, as further described below, reveals that different types of interactions govern these transitions (Fig. 4b). In general, water-mediated interactions primarily drive changes at the intradimer interfaces, highlighting their role in modulating lattice dynamics. Conversely, direct contact changes are predominant at the interdimer interface. Changes at lateral interfaces typically involve both direct and water-mediated contacts.

The initial transition to the compacted state (from GMPCPP-to GDP·BeF₃⁻) results in a net increase in both intradimer and interdimer longitudinal contacts (Fig. 4bc, Fig. S7ab). Structural rearrangements promote new interdimer interactions between βT5, βH5, and βH6, and between αH10 and αH10–αS9, consistent with previous studies (*34*). At the same time, we observed an increased number of contacts between αT7 and αH8 with the nucleotide, which is consistent with activation of the catalytic site. In contrast, lateral contacts are diminished during compaction (Fig. 4b, Fig. S7c), underscoring their role in stabilizing the expanded GTP-lattice (*19*).

The subsequent hydrolysis-associated transition (from GDP·BeF₃⁻ to GDP·AlF₃) elicits mild but opposite responses at the two longitudinal interfaces. The intradimer interface experiences a net loss of contacts, whereas the interdimer interface exhibits a net gain (Fig. 4bd, Fig. S7ab). Together with a net gain in contacts at the α–α’ lateral interface (Fig. 4b, Fig. S7c), these conformational effects increase the connectivity between tubulin dimers, which may help maintain lattice integrity during hydrolytic cleavage of the β–γ phosphate ester bond during this transition. Additionally, βT3 forms extensive interactions with αH8 and the catalytic loop αT7 (Fig. S7a), providing structural evidence for the previous proposal that it acts as a sensor of the nucleotide state (*34, 35*).

Lastly, the transition to the GDP state (from GDP·AlF₃ to GDP) represents the release of Pi and the Mg²⁺ ion. This terminal event triggers a marked loss of contacts at both the longitudinal interdimer and the lateral interfaces (Fig. 4be, Fig. S7). The intradimer interface undergoes no further changes, effectively preserving the relative instability acquired during the preceding steps of GTP hydrolysis and priming the GDP-microtubule lattice for rapid disassembly.

### Conclusions

Our findings establish a detailed structural framework linking the chemical steps of GTP hydrolysis to the regulation of microtubule lattice dynamics. Crucially, our data identify ordered water molecules as essential components of the lattice interaction network. Together with protein-protein interactions, water molecules link GTP hydrolysis at the catalytic site to large-scale conformational transitions that occur throughout the microtubule lattice and involve both intradimer and interdimer contacts.

In this framework, the transition from an expanded to a compacted GTP lattice is a pre-requisite for efficient GTP hydrolysis and emerges as a crucial mechanistic checkpoint. While an expanded GTP cap is a well-established characteristic of growing microtubule ends (*20, 21*) and is likely essential for lattice elongation and closure (*36, 37*), our findings indicate that transitioning to the compact conformation is necessary to facilitate hydrolysis. This pre-hydrolysis compaction mechanism carries several important implications. First, by optimizing the arrangement of catalytic residues, the Mg²⁺ ion, and ordered water molecules within the catalytic site, lattice compaction can explain the nearly thousand-fold increase in the rate of GTP hydrolysis observed when tubulin is incorporated into microtubules (*14*). Second, the requirement for lattice compaction to promote GTP hydrolysis provides insight into why GTP hydrolysis is delayed relative to tubulin incorporation at growing microtubule ends (*1, 3, 10*). Finally, the pre-hydrolysis compaction mechanism offers a structural explanation for the GAP-like activity of tubulin within the microtubule lattice postulated more than 30 years ago (*15–17*).

Another key insight from our work is that local GTP hydrolysis within the catalytic site exerts a global influence on lattice-incorporated tubulin dimers. The transition from the compact GTP (GDP·BeF₃⁻) to the GDP·Pᵢ (GDP·AlF₃) microtubule lattice state involves a widespread redistribution of lattice interactions that maintain lattice integrity during hydrolytic cleavage of the γ-phosphate. These conformational changes exemplify the coupling between an atomic-scale event at the catalytic site, namely γ-phosphate cleavage, and large-scale remodeling of the microtubule lattice. The subsequent transition from the GDP·Pᵢ to the final GDP state further highlights this mechanistic framework: the release of Pᵢ and Mg²⁺ together with the remodeling of the catalytic site are atomic-level events that drive a substantial net loss of lattice contacts. This finding provides a structural basis for understanding how lattice strain promotes microtubule destabilization and ultimately triggers catastrophe, a hallmark of microtubule dynamic instability.

Together, our findings provide a coherent structural basis linking nucleotide state, lattice conformation, and microtubule stability throughout the GTP hydrolysis reaction. More broadly, the ability to resolve microtubule structures at increasingly high resolutions, combined with the structural insights obtained here, establish a foundation for future studies of microtubule biology at the molecular level, including investigations into how disease-associated tubulin mutations, microtubule-associated proteins, and microtubule drugs modulate microtubule lattice structure and dynamics.

## Materials and Methods

### Tubulin and microtubule preparations

Lyophilized bovine brain tubulin (Centro de Investigaciones Biológicas Margarita Salas, CSIC, Madrid, Spain) was resuspended in BRB80 buffer (80 mM PIPES, 1 mM EGTA, 1 mM MgCl₂, pH 6.8) supplemented with 0.1 mM GTP (NU-1047, Jena Bioscience) to a final concentration of 10 mg/mL and subjected to one cycle of polymerization–depolymerization, based on a previously described protocol (*38*). Briefly, the tubulin solution was supplemented with 3.4 M glycerol (104091, Merck-Sigma Aldrich) and 1 mM GTP. Mixtures were incubated at 37 °C for 45 min to induce microtubule polymerization. Polymerized microtubules were pelleted by centrifugation at 16,000 × g for 20 min at 37 °C. The pellet was resuspended in ice-cold BRB80 supplemented with 0.1 mM GTP and homogenized gently. The suspension was then clarified by ultracentrifugation at 80,000 × g for 15 min at 4 °C to remove aggregates and insoluble material. The supernatant containing soluble tubulin was collected, and protein concentration was determined using a NanoDrop spectrophotometer (Thermo Fischer Scientific). Aliquots were snap-frozen in liquid nitrogen and stored at −80 °C until use.

The following microtubule samples were prepared:

#### GMPCPP-microtubules

Preparation of GMPCPP-microtubules was performed based on a previous protocol (*23*). Briefly, 25 μL of cycled tubulin was diluted to 2 mg/mL in BRB80 supplemented with 0.1 mM GTP. This tubulin solution was supplemented with 1 mM GMPCPP (NU-405, Jena Bioscience) and incubated at 37 °C for 1 h to allow polymerization, followed by centrifugation at 16,000 × g for 20 min at 37 °C. The resulting pellet was resuspended in 50 μL of warm BRB80 containing 0.1 mM GMPCPP. Samples were maintained at room temperature and used within 30 min for grid preparation.

#### GDP·BeF₃⁻-microtubules

First, a BeF₃⁻-stock solution was prepared as previously described (*24*). A 55.6mM stock was prepared by mixing 1 molar equivalent of BeSO₄ (14270, Merck-Sigma Aldrich) and 3 molar equivalents of HKF_2_ (239283, Merck-Sigma Aldrich) in Milli-Q water and adjusted to pH 5–6. GDP·BeF₃⁻-microtubules were assembled using a self-seeding protocol (*39*). First, GDP·BeF₃⁻-microtubule seeds were generated by diluting 25 μL of cycled tubulin to 4 mg/mL in BRB80 supplemented with 3.4 M glycerol and 10 mM BeF₃⁻. The mixture was incubated at 37 °C for 1 h and centrifuged at 16,000 × g for 20 min at 37 °C. Pellets were resuspended in 25 μL of warm BRB80 supplemented with 10 mM BeF₃⁻ and subjected to two rounds of 30 s sonication, with vigorous pipetting between cycles to generate short seeds. Final GDP·BeF₃⁻-microtubules were obtained by adding 10% (v/v) of the seed suspension to tubulin at 5 mg/mL in BRB80 (without GTP) supplemented with 10 mM BeF₃⁻, followed by incubation at 37 °C for at least 1 h.

#### GDP·AlF₃-microtubules

GDP·AlF₃-microtubules were prepared as previously described (*24, 39*). Briefly, GDP·AlF₃-microtubule seeds were generated by diluting 25 μL of cycled tubulin to 7 mg/mL in BRB80 supplemented with 3.4 M glycerol, 2.5 mM GDP (NU-1172, Jena Bioscience), 1.5 mM AlCl₃ (1.28223, Merck-Sigma Aldrich), and 3 mM HKF_2_, allowing in situ formation of AlF₃. The mixture was incubated at 37 °C for 1 h and centrifuged at 16,000 × g for 20 min at 37 °C. Pellets were resuspended in BRB80 containing 1.5 mM AlCl₃, 3 mM HKF_2_, and 2.5 mM GDP Seed preparation by sonication and microtubule elongation were performed as described above for BeF₃⁻ samples, using polymerization buffer supplemented with 1.5 mM AlCl₃, 3 mM HKF_2_, and 1 mM GDP.

#### GDP-microtubules

To assemble GDP microtubules, 25 μL of cycled tubulin at 8 mg/ml in BRB80 were supplemented with 10% glycerol and 2 mM GTP. The mixture was then incubated at 37 °C for 2 h to allow polymerization, followed by centrifugation at 16,000 × g for 20 min at 37 °C. The resulting pellet was resuspended in 50 μL of warm BRB80 containing 10% glycerol and 2 mM GTP. Samples were maintained at 37 °C and used within 30 minutes for grid preparation.

### Cryo-EM grid preparation

Cryo-EM grids were prepared using Quantifoil R 2/1 holey carbon films on 300-mesh copper grids (QuantiFoil Micro Tools, GmbH). Prior to sample application, grids were glow-discharged for 30 s at 20 mA, negative polarity, using a GloQube Plus glow discharge system (Quorum Technologies). GDP·BeF₃⁻- and GDP-microtubule samples were vitrified using a Vitrobot Mark IV (Thermo Fisher Scientific), applying a blot force of 22 (instrument units). GMPCPP- and GDP·AlF₃-microtubules were vitrified using an EM GP2 (Leica Microsystems). For GDP microtubules, the environmental chamber was equilibrated at 37 °C and 100% relative humidity. 2.5 μL of microtubule sample was applied to each grid and blotted for 12 s. For all other conditions, the environmental chamber was equilibrated at 30 °C and 90% relative humidity prior to vitrification. A volume of 3 μL of microtubule sample was applied to each grid and blotted immediately using the following conditions: 4 s for GMPCPP and GDP·AlF₃ samples, and 6 s for GDP·BeF₃⁻ samples. Grids were plunge-frozen in liquid ethane immediately after blotting and subsequently transferred to liquid nitrogen for storage. All grids were stored under liquid nitrogen and used for data acquisition within two weeks of preparation.

### Cryo-EM data collection and processing

Prior to data acquisition, grids were screened for ice thickness, contamination, and filament distribution using a Talos F200C (Thermo Fisher Scientific) operated at 200kV and equipped with a Ceta 16M detector (Thermo Fischer Scientific). Grid squares exhibiting optimal ice quality and appropriate filament density were selected for high-resolution data collection using a Titan Krios G4 (Thermo Fisher Scientific) operated at 300 kV, equipped with a Falcon 4i direct electron detector and a Selectris X energy filter (Thermo Fischer Scientific). Data collection parameters are detailed in Table S1.

Cryo-EM data was processed using our MiCSPARC pipeline (*40*) including CryoSPARC (*41*) (v4.7.1 and 5.0.4). Briefly, micrograph-movies were motion-corrected patch-wise using default settings. The micrographs’ CTFs were estimated patch-wise, and those where the CTF was fitted to a resolution better than 4.5 Å were included in further processing. Helical particles were picked from filaments with 82 Å spacing and 2D-classified to select microtubule particles. The resulting datasets were extrapolated using MiCSPARC script to ensure correct filament assignments. Extrapolated particles were class assigned by 3D heterogeneous refinement using 6 generated references with theoretical helical parameters for 11-3, 12-3, 13-3, 14-3, 15-4, and 16-4 microtubules. The class assignments were then probabilistically smoothed using MiCSPARC script. The largest resulting classes (GDP 13-3 and 14-3, GDP·BeF_3_^-^ 13-3, GDP·AlF_3_ 13-3 and GMPCPP 13-3 and 14-3) were further processed.

Selected particles were submitted to an initial helix refinement with enforced symmetry parameters, to produce a roughly aligned particle set and an appropriate helical mask. Then, particle angles were unified using MiCSPARC. First, in-plane rotation was unified for each filament. Then, two iterations of phi angle unification were performed. The resulting particle set was symmetry-expanded and refined without symmetry. A mask for a single pf was prepared, and the particles and volume were shifted to center on that pf. New, smaller particles were extracted from micrographs at every symmetry-related position in the dataset and reconstructed into a pf structure.

To distinguish α- and β-tubulin monomers in our pf reconstructions, the pf particles were classified into two classes of different registers, using S9-S10 loops on the luminal side of the filament as reference. One of the classes was shifted 41 Å along the helical axis, and both classes were locally refined together.

The defoci of the particles and higher-order CTF aberrations were estimated and refined. The part of the structure outside of the central three tubulin monomers was subtracted, and the subtracted particles were locally refined. The final pf structure, containing all the symmetry-expanded particles, was then subject to particle filtering by high-resolution 3D classification, filtering on scale factor and out-of-plane tilt. Local resolution and local filtering in CryoSPARC produced the final tubulin trimer maps that were used for model building.

The seam position of the microtubules was determined by MiCSPARC from the final full particle set and the initial register assignment of 3D classification, assuming all microtubules have a single seam. For each particle, probability that it is placed at the seam was calculated from the register assignment of all its neighbors within each symmetry-expansion set. The particles most likely to be located at the seam were shifted to center the particle on the microtubule axis and re-extracted to reconstruct a full microtubule volume. The particle set was then locally refined to produce the fully seamed microtubule volume, used to determine the lattice parameters.

### Model building and refinement

Atomic models were built in Coot (*42*) (v0.9.8.95) using local filtered maps. We used minimized pf maps in which an αβ-tubulin heterodimer (α-tubulin, chain A; β-tubulin, chain B) and an additional β-tubulin monomer from another, longitudinally aligned tubulin dimer (chain C) are reconstructed (Fig. S2). Initially, a minimized pf model (*43*) (PDB: 6WVR) was rigid-body fitted into cryo-EM density maps using the jiggle-fit tool. An initial round of water addition was performed in Phenix (*44*) (v1.21.2.5419), using phenix.douse. Macromolecular geometry, secondary structures, backbone and lateral chain, ligands, and waters were manually adjusted into the density of each map. A model to map refinement was performed in Phenix using the phenix.real_space_refine program. Iterative cycles of model building and refinement were performed until good stereochemistry and agreement of the models with the electron density were achieved, with optimal refinement statistics. Waters were modelled only when supported by the density contoured at > 1.0 RSMD and a chemically reasonable local environment. Water molecules were further validated by Q-score analysis (*27*) (Fig. S9). Other solvent molecules were modelled only when supported by density contoured at > 1.0 RMSD. None of these induced detectable changes in the local protein environment.

### Model analysis

Structural models were analyzed using ChimeraX (*45*) v1.10.1 and python libraries numpy, pandas, and matplotlib (*46–48*). All figures were prepared using ChimeraX, matplotlib, and Inkscape v1.4.2 (The Inkscape Project, Inkscape (Version 1.4.2), https://inkscape.org).

#### Catalytic pocket interaction networks

The highest resolution models from each microtubule structure (14-pf GMPCPP, 13-pf GDP·BeF_3_^-^, 13-pf GDP·AlF_3_, 14-pf GDP) were aligned on all the matching atoms of the nucleotide present in β_2_-tubulin, and the angles and distances were measured using ChimeraX. Eight bond angles describing the Mg^2+^coordination were measured, and their standard deviation calculated. Six distances from the Mg^2+^ cation to each coordinating atom were measured, and their standard deviation calculated. Hydrogen bonds were calculated using ChimeraX based on default criteria (*49*).

#### Lattice parameters

Intradimer distance, interdimer distance, and dimer rise were calculated from fitted 13-pf models as average pairwise distance between corresponding C_α_ atoms (*22*). The pf twist was measured from seamed microtubule volumes using CryoSPARC. Tubulin backbone movements were analyzed in two superpositions: the movements inside the tubulin dimer α_1_β_1_ were visualized on structures aligned on the N-terminal domain (residues 1-205) of α_1_-tubulin. The movements between two tubulin dimers were visualized on structures aligned on the N-terminal domain (residues 1-205) of β_2_-tubulin. The direction of movement of every C_α_ in each structural transition was plotted as described previously (*50*) and for clarity in the figures, the magnitude of movement is shown 5-fold larger.

#### Contact maps

Hexamer models containing two tubulin trimers in adjacent pfs were prepared by rigid-body fitting the 13-pf trimer models into a pre-particle subtraction pf map. Clashing waters were removed from the models on lateral interfaces. The AlF_3_ and BeF_3_^-^ atoms were assigned to chemical identities of O instead of F, and P instead of Be and Al. Hydrogen bond list was generated using ChimeraX with tolerance increased by 0.1 Å and 10°. Atom-atom contact list was generated using ChimeraX with default settings. These lists were analyzed using numpy and pandas in python. Polar contacts for each interface were defined as direct hydrogen bond interactions between two tubulin chains, treating the associated nucleotide, BeF_3_^-^ and AlF_3_ moieties as part of the protein chain. Water-mediated interactions were defined as a set of waters that simultaneously form at least 1 hydrogen bond to each neighboring protein chain. Water bridges requiring more than one water were not included in this analysis. Hydrophobic contacts were defined as carbon-carbon contacts within a distance suitable for van der Waals interaction, counting only a single such interaction per residue pair. Secondary structure assignments were mapped according to ref. (*28*). Contact maps were prepared using matplotlib and Inkscape.

## Acknowledgments

We thank Dr. Rémi Ruedas and Dr. Mohammed Chami from the BioEM lab of the Biozentrum at the University of Basel for excellent support with cryo-EM grid screening and data collection. We are indebted to Dr. Andrea Prota and Dr. Daniel Lucena-Agell for critical reading of the manuscript.

## Author contributions

**J.E.-G**. conceptualized and designed the research, prepared microtubule samples and cryo-EM grids, supervised the cryo-EM data collection, processed cryo-EM data, performed model building and validation, analyzed structures, and wrote the manuscript. **P.F.** conceptualized and designed the research, prepared microtubule samples and cryo-EM grids, supervised the cryo-EM data collection, processed cryo-EM data, performed model building and validation, analyzed structures, prepared figures, and contributed to the writing of the manuscript. **H.M.-H**. performed model building and validation. **S.M.** analyzed structures, validated models, and contributed to figure preparation. **M.W.** directed the project, conceptualized the research, supervised the model building and structure analysis, and contributed to the writing of the manuscript and to figure preparation. **M.O.S.** directed the project, conceptualized and designed the research, supervised the structure analysis, and wrote the manuscript.

## Data availability

The atomic model coordinates and related cryo-EM maps of each protofilament structure presented in this work have been deposited at the Protein Data Bank (PDB) with the following accession numbers: 31CP (13-pf GMPCPP-microtubules), 31CQ (14-pf GMPCPP-microtubules), 31CL (13-pf GDP·BeF3—microtubules), 31CM (13-pf GDP·AlF3-microtubules), 31CN (13-pf GDP-microtubules), and 31CO (14-pf GDP-microtubules). Associated maps related to the microtubule reconstruction with the seam search approach following the MiCSPARC (*40*) pipeline were also deposited with the following accession numbers: EMD-58297 (13-pf GMPCPP-microtubules), EMD-58298 (14-pf GMPCPP-microtubules), EMD-58292 (13-pf GDP·BeF3—microtubules), EMD-58293 (13-pf GDP·AlF3-microtubules), EMD-58295 (13-pf GDP-microtubules), and EMD-58296 (14-pf GDP-microtubules).

## Declaration of generative AI and AI-assisted technologies in the writing process

During the preparation of the manuscript, the authors used ChatGPT (OpenAI, LLC), Gemini 3.1 Pro (Alphabet Inc.) and Perplexity (Perplexity AI, Inc.) only to improve the language. After using this tool, the authors reviewed and edited the content as needed, and they take full responsibility for the content of the publication.

