## Supplementary materials for "Structural basis of microtubule destabilization by GTP hydrolysis"

**This document includes:**

- Tables S1 to S5
- Figures S1 to S9
- Legends Movie S1 and S2
- Supplementary References

### Supplementary Tables

#### Table S1. Cryo-EM data collection.

| | GMPCPP | GDP·$BeF_3^-$ | GDP·$AlF_3$ | GDP |
|---|---|---|---|---|
| **Voltage (keV)** | 300 | 300 | 300 | 300 |
| **Pixel size (Å/px)** | 0.738 | 0.738 | 0.738 | 0.946 |
| **Camera** | Falcon 4i | Falcon 4i | Falcon 4i | Falcon 4i |
| **Acquisition Mode** | Counting | Counting | Counting | Counting |
| **Total dose (e/Å$^2$)** | 30 | 30 | 30 | 30 |
| **Energy filter slit width (eV)** | 10 | 10 | 10 | 10 |
| **Defocus range** | −0.8 to −2.0 | −0.8 to −2.0 | −0.8 to −2.0 | −0.6 to −1.8 |
| **Number of movies** | 8,000 | 4,246 | 11,193 | 8,438 |

### Table S2. Cryo-EM data processing.

| | GMPCPP | GMPCPP | GDP·BeF3- | GDP·AlF3 | GDP | GDP |
|---|---|---|---|---|---|---|
| **Protofilament number** | 13 | 14 | 13 | 13 | 13 | 14 |
| **Processing pixel size (Å/pixel)** | 0.738 | 0.738 | 0.738 | 0.738 | 0.946 | 0.946 |
| **Box size (pixels)** | 360 | 360 | 360 | 360 | 320 | 256 |
| **Extracted segments** | 174,551 | 174,551 | 87,301 | 186,268 | 100,441 | 100,441 |
| **Resolution (Å)** | 2.13 | 1.88 | 2.02 | 1.97 | 2.17 | 2.04 |
| **Number of particles** | 219,377 | 1,334,721 | 352,525 | 413,274 | 349,895 | 785,248 |
| **Accession numbers** | PDB-31CP<br>EMD-58297 | PDB-31CQ<br>EMD-58298 | PDB-31CL<br>EMD-58292 | PDB-31CM<br>EMD-58293 | PDB-31CN<br>EMD-58295 | PDB-31CO<br>EMD-58296 |

**Table S3. Seam-corrected microtubule reconstruction details**

| | GMPCPP | GDP·BeF3- | GDP·AlF3 | GDP | GMPCPP | GDP |
|---|---|---|---|---|---|---|
| **Protofilament number** | 13 | 13 | 13 | 13 | 14 | 14 |
| **Number of particles** | 19,437 | 59,216 | 51,434 | 39,427 | 133,917 | 43,812 |
| **Resolution (Å, FSC 0.143)** | 3.49 | 3.20 | 3.03 | 3.27 | 2.85 | 3.11 |
| **Seam identification** | Yes | Yes | Yes | Yes | Yes | Yes |
| **Model building** | No | No | No | No | No | No |
| **Accession number** | EMD-58275 | EMD-58204 | EMD-58234 | EMD-58218 | EMD-58276 | EMD-58294 |

### Table S4. Tubulin trimer model building and refinement.

| | GMPCPP | GMPCPP | GDP·BeF3- | GDP·AlF3 | GDP | GDP |
|---|---|---|---|---|---|---|
| Pf number | 13 | 14 | 13 | 13 | 13 | 14 |
| Initial model used | 6WVR | | | | | |
| $FSC_{model, masked}$ (0/0.143/0.5) (Å) | 1.8/2.1/2.3 | 1.5/1.7/1.9 | 1.7/1.9/2.1 | 1.6/1.8/2.0 | 1.8/2.0/2.2 | 1.8/2.0/2.1 |
| $CC_{mask}$ | 0.78 | 0.90 | 0.89 | 0.89 | 0.86 | 0.88 |
| **Model composition** | | | | | | |
| Atoms (hydrogens) | 20643 (9852) | 20980 (9965) | 20941 (9870) | 21064 (9945) | 20771 (9907) | 20929 (9901) |
| Protein residues | 1,282 | 1,284 | 1,291 | 1,291 | 1,290 | 1,291 |
| Ligands | 6 | 6 | 8 | 10 | 4 | 4 |
| Waters | 568 | 674 | 796 | 779 | 575 | 754 |
| **B factors (Å$^2$)** | | | | | | |
| Protein (min/max/avg) | 9.01/84.30/32.66 | 2.91/60.85/18.17 | 2.84/67.39/22.66 | 0.83/78.59/16.86 | 7.38/73.19/29.71 | 5.97/78.28/24.75 |
| Ligand (min/max/avg) | 14.68/43.98/26.55 | 4.54/28.03/12.64 | 2.77/31.93/29.67 | 2.70/67.88/28.25 | 13.03/28.40/19.01 | 8.58/26.43/14.83 |
| Water (min/max/avg) | 12.86/63.31/37.31 | 6.20/44.12/26.54 | 4.44/50.62/29.67 | 2.20/53.31/25.65 | 14.62/53.41/35.38 | 8.89/57.48/33.74 |
| **r.m.s. deviations** | | | | | | |
| Bond lengths (Å) | 0.003 | 0.004 | 0.005 | 0.004 | 0.003 | 0.003 |
| Bond angles (°) | 0.544 | 0.597 | 0.615 | 0.591 | 0.521 | 0.55 |
| **Validation** | | | | | | |
| MolProbity score | 1.3 | 0.89 | 0.83 | 0.96 | 0.8 | 0.84 |
| Clashscore, all atoms | 2.09 | 1.52 | 1.39 | 1.93 | 1.04 | 1.24 |
| Cβ outliers (%) | 0 | 0 | 0 | 0 | 0 | 0 |
| CaBLAM outliers (%) | 0.47 | 0.47 | 0.39 | 0.78 | 0.39 | 0.47 |
| Poor rotamers (%) | 1.81 | 0.71 | 0.81 | 0.54 | 0.99 | 0.45 |
| **Ramachandran plot** | | | | | | |
| Favored (%) | 97.33 | 98.75 | 98.75 | 98.29 | 98.52 | 98.36 |
| Allowed (%) | 2.67 | 1.25 | 1.25 | 1.71 | 1.64 | 1.78 |
| Disallowed (%) | 0 | 0 | 0 | 0 | 0 | 0 |
| **Rama-Z score** | | | | | | |
| Whole | 0.33 | 0.33 | 0.42 | 0.43 | 0.7 | 0.56 |
| Helix | 0.64 | 0.81 | 0.78 | 0.96 | 1.68 | 0.99 |
| Sheet | -0.35 | -0.18 | 0.15 | 0.03 | 0.3 | 0.34 |
| Loop | -0.36 | -0.08 | -0.05 | -0.23 | -0.83 | -0.18 |

### Table S5. Microtubule lattice parameters.

| | GMPCPP | GDP·BeF3- | GDP·AlF3 | GDP | GMPCPP | GDP |
|---|---|---|---|---|---|---|
| **Protofilament number** | 13 | 13 | 13 | 13 | 14 | 14 |
| **Average dimer rise [Å]** | 84.88 | 82.27 | 82.30 | 82.44 | 84.87 | 82.43 |
| **Volume dimer twist [°]** | 0.17 | -0.07 | -0.08 | 0.04 | -0.46 | -0.45 |
| **Average intradimer distance [Å]** | 41.79 | 41.73 | 41.68 | 41.78 | 41.79 | 41.94 |
| **Average interdimer distance [Å]** | 43.10 | 40.53 | 40.66 | 40.70 | 43.12 | 40.52 |

### Supplementary Figures

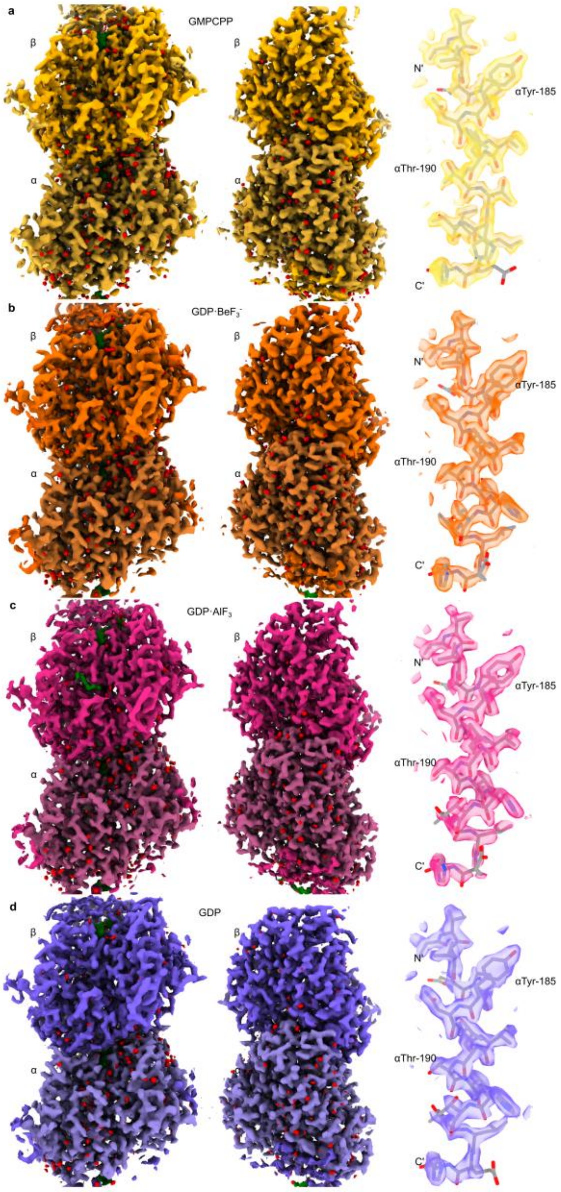


**Figure S1. Sub-2.2 Å resolution microtubule structures.** Luminal (left) and cytoplasmatic (center) views of cryo-EM density maps of (**a**) 14-pf GMPCPP-microtubule at 1.9 Å (yellow). (**b**) 13-pf GDP·$BeF_3^-$-microtubule at 2.02 Å (orange). (**c**) 13-pf GDP·$AlF_3$-microtubule at 2.0 Å (magenta). (**d**) 14-pf GDP-microtubule at 2.0 Å (purple) contoured at 1.5 RMSD. Ligands and waters are colored in green and red, respectively. The right panel contains close-up views of helix αH5, with the corresponding models (stick representation) and density maps contoured at 2.5 RMSD.

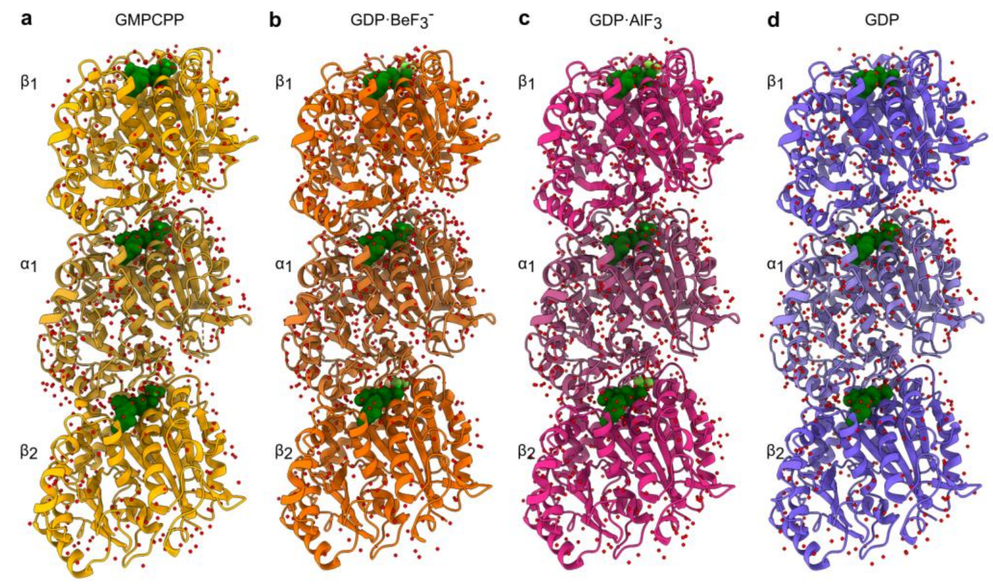


**Figure S2. Microtubule trimer models.** Cartoon representation of the luminal view of trimer models. Each model is composed of a tubulin dimer ($\alpha_1$-$\beta_1$) and an additional beta subunit ($\beta_2$). Panels correspond to different nucleotide states: (**a**) 14-pf GMPCPP-microtubule (yellow). (**b**) 13-pf GDP·$BeF_3^-$ -microtubule (orange). (**c**) 13-pf GDP·$AlF_3$-microtubule (magenta). (**d**) 14-pf GDP-microtubule (purple). Waters and $Mg^{2+}$ ions are represented as red and lime spheres, respectively. The nucleotides are shown in green spheres representation.

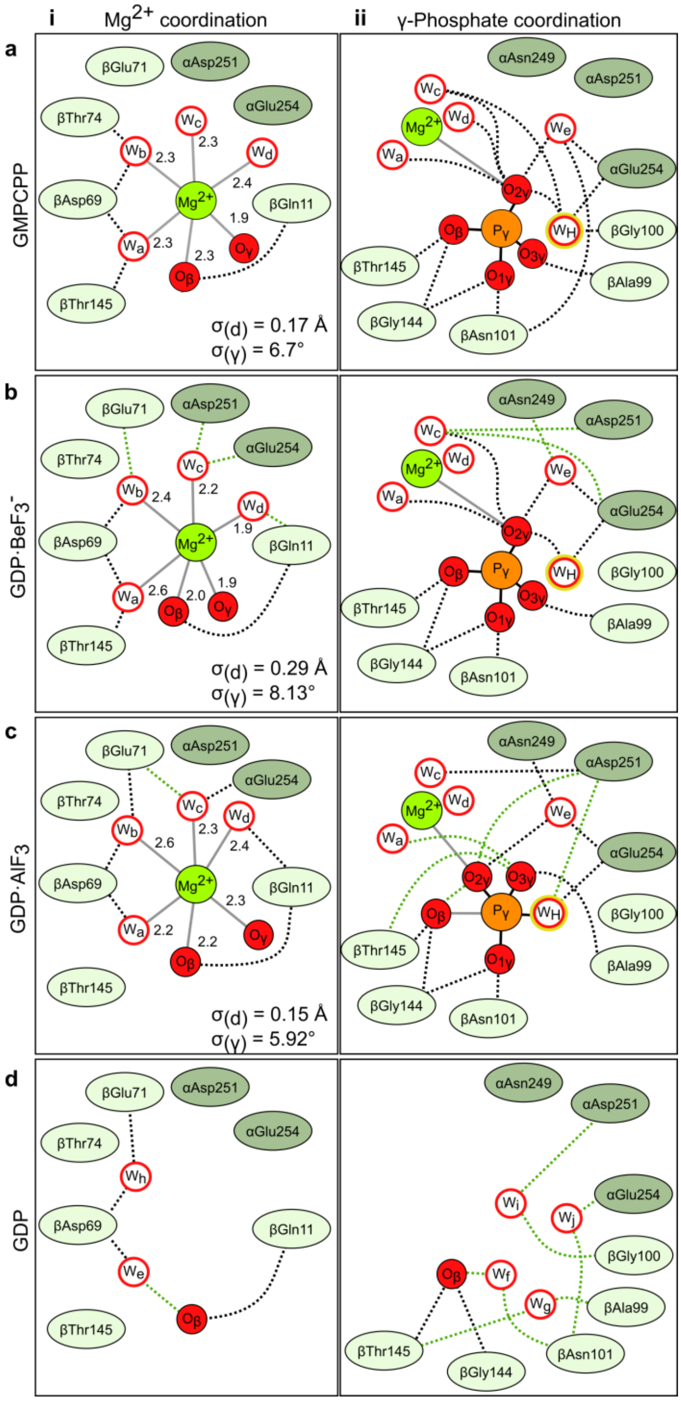


**Figure S3. $Mg^{2+}$ ion and γ-phosphate interaction networks at the catalytic site.** Schematic representation of the interaction networks of the $Mg^{2+}$-water coordination sphere (**i**) and the nucleotide γ-phosphate (**ii**) for GMPCPP- (**a**), GDP·$BeF_3$- (**b**) and GDP·$AlF_3$-microtubules (**c**). The empty pocket in the GDP-microtubule is represented in (**d**). Distances are illustrated as grey lines and reported in Å. The standard deviation of the $Mg^{2+}$ coordination bond lengths ($\sigma_d$) and angles ($\sigma_\gamma$) are given. Water molecules are represented as white circles with a red rim, phosphate oxygens, phosphorus, and $Mg^{2+}$ ion as solid circles in red, orange, and green, respectively. Residues in the pocket are shown in light green for β-tubulin and dark green for α-tubulin. Covalent bonds are shown as solid lines. Conserved hydrogen bonds throughout the transitions are shown as black dashed lines. Newly formed hydrogen bonds are represented as green dashed lines. The catalytic water is highlighted with a yellow rim.

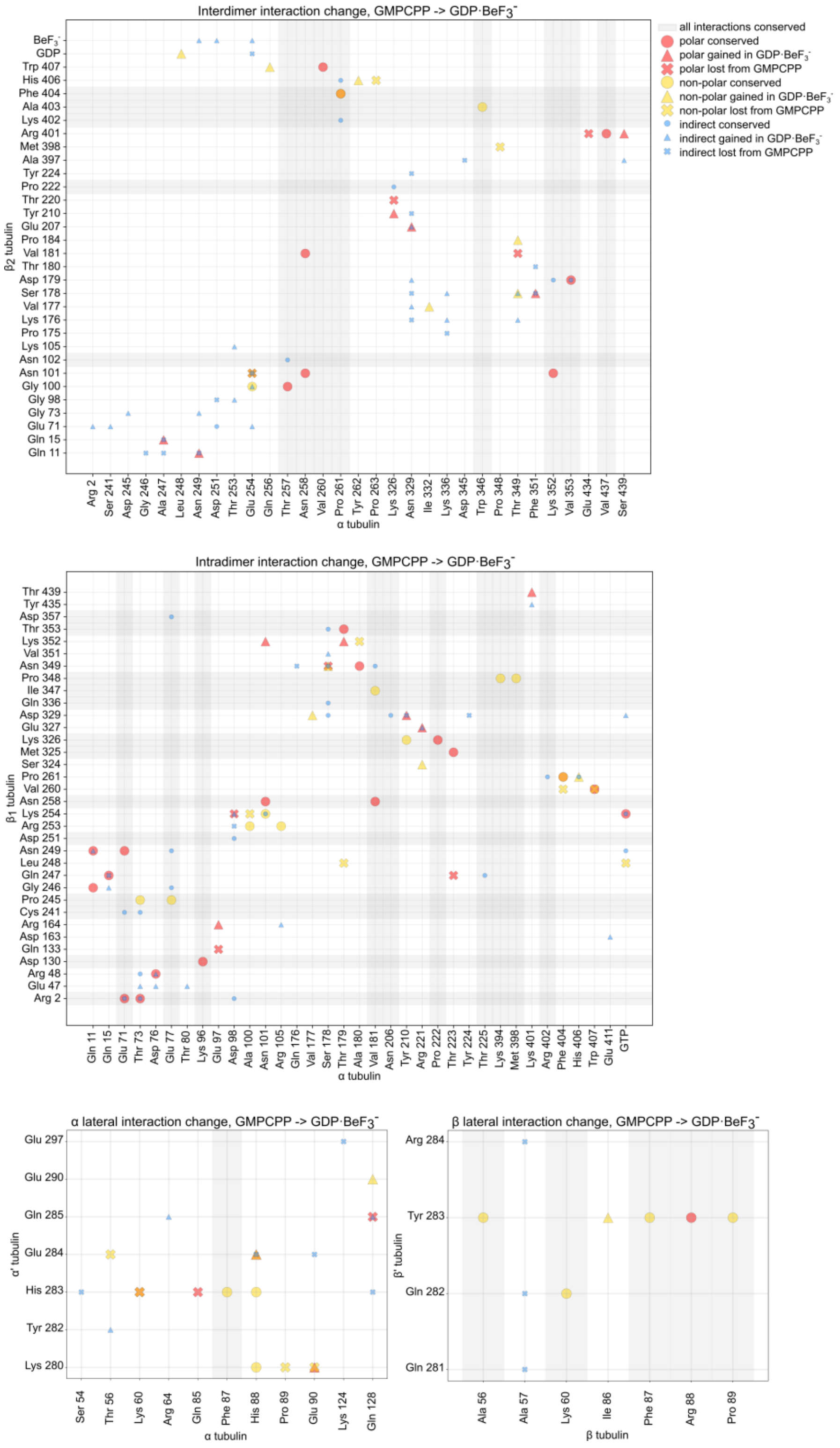


**Figure S4. Interaction plots for the GMPCPP- to GDP·$BeF_3^-$-microtubule lattice transition.** Plots depict changes in interactions at the Interdimer (top), Intradimer (center) α–α’ (bottom left) and β–β’ interfaces (bottom right) upon the GMPCPP to GDP·$BeF_3^-$ microtubule transition. The x- and y-axes represent the α- and β-tubulin residues involved in the interaction changes, respectively. Symbols are defined in the key legend (top, right). Gray lines indicate zero changes in the number of contacts.

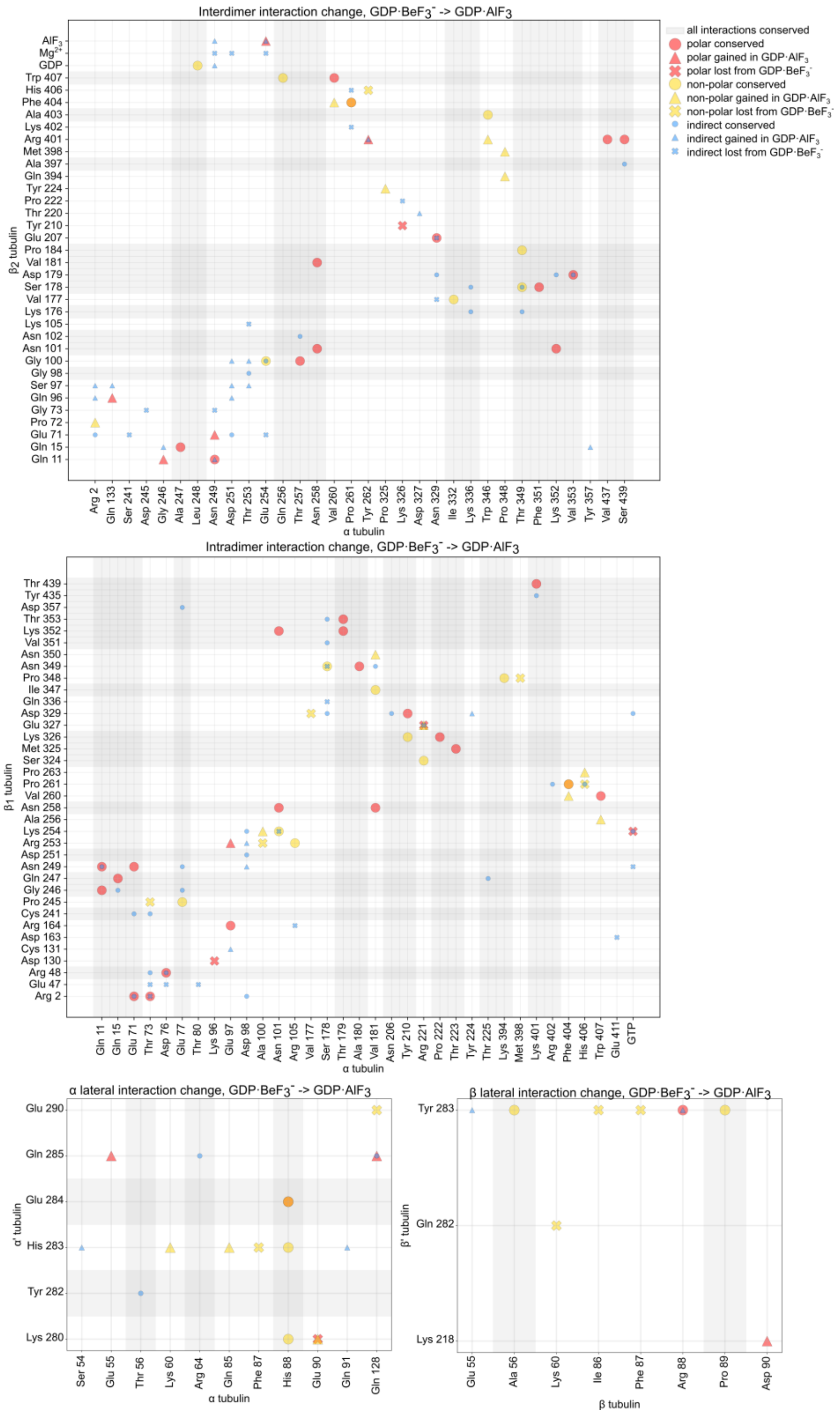


**Figure S5. Interaction plots for the GDP·BeF$_3^-$- to GDP·AlF$_3$-microtubule lattice transition.** Plots depict changes in interactions at the Interdimer (top), Intradimer (center) α–α' (bottom left) and β–β' interfaces (bottom right) upon the GDP·BeF$_3^-$ to GDP·AlF$_3$ microtubule transition. The x- and y-axes represent the α- and β-tubulin residues involved in the interaction changes, respectively. Symbols are defined in the key legend (top, right). Gray lines indicate zero changes in the number of contacts.

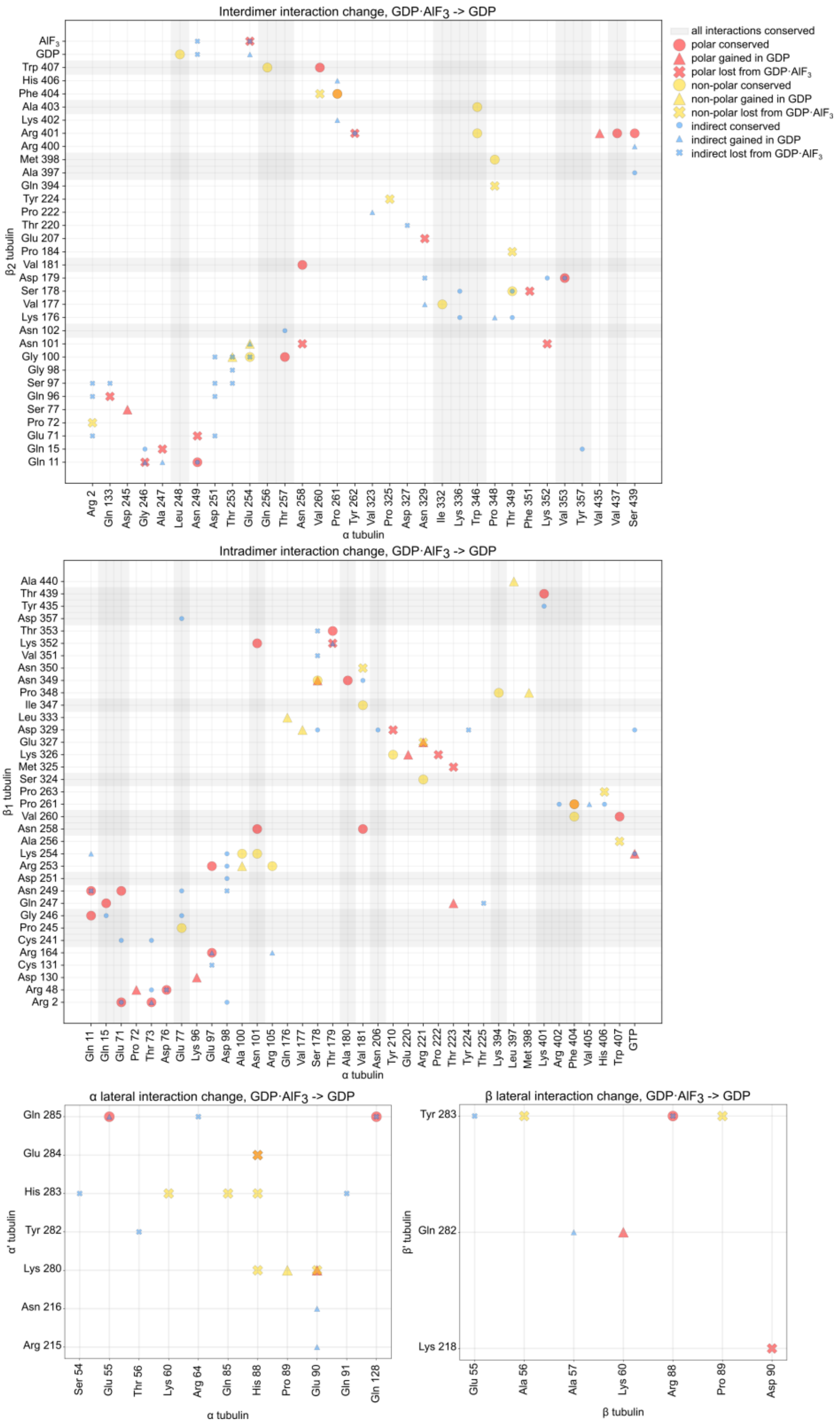


**Figure S6. Interaction plots for the GDP·BeF$_3^-$- to GDP-microtubule lattice transition.** Plots depict changes in interactions at the Interdimer (top), Intradimer (center) α–α (bottom left) and β–β interfaces (bottom right) upon the GDP-BeF$_3^-$ to GDP microtubule transition. The x- and y-axes represent the α- and β-tubulin residues involved in the interaction changes, respectively. Symbols are defined in the inset legend (top right). Symbols are defined in the key legend (top, right). Gray lines indicate zero changes in the number of contacts.

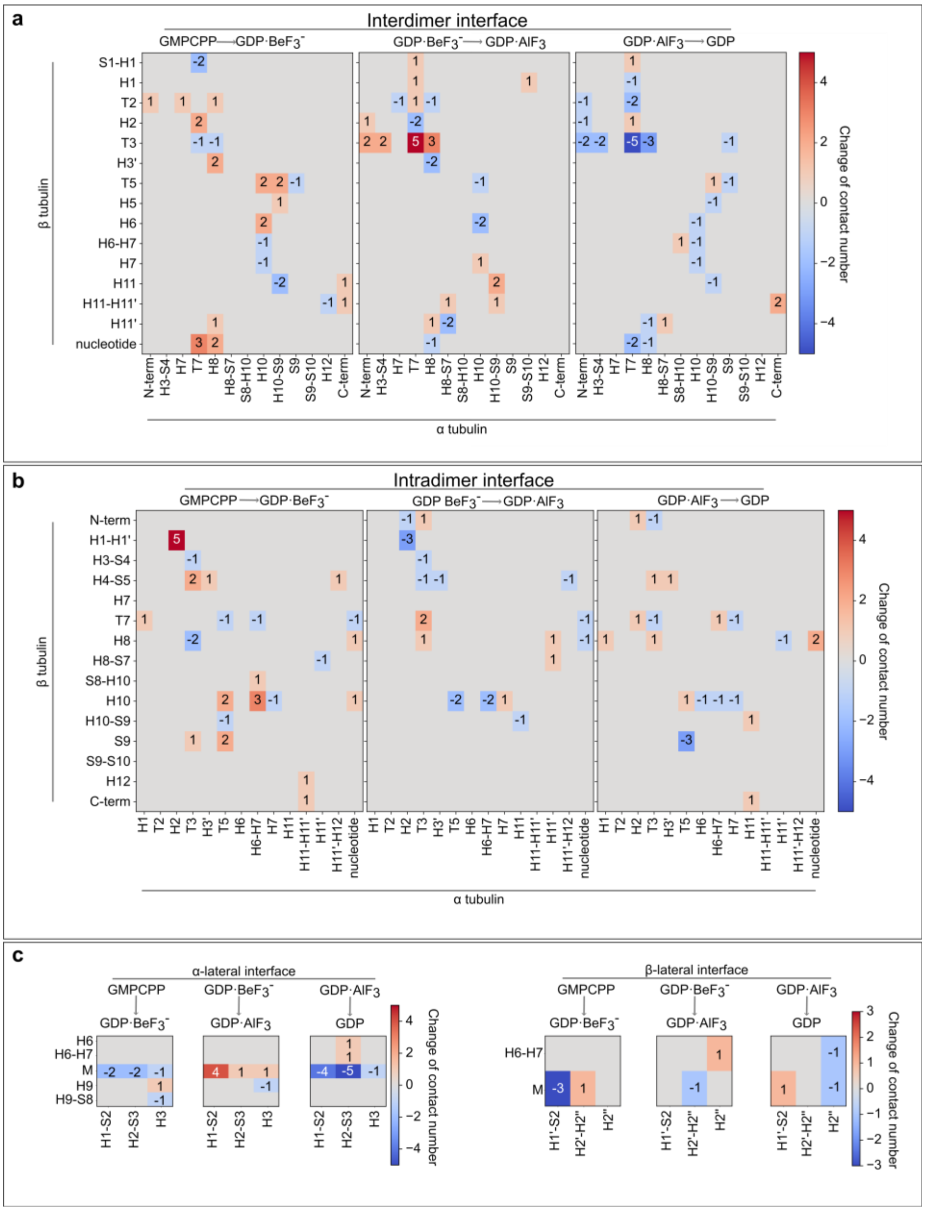


**Figure S7. Secondary structure contact changes throughout the GTP hydrolysis reaction.** Heatmaps describing local changes in contact numbers at the longitudinal interdimer (**a**), longitudinal intradimer (**b**), and lateral α–α' (**c, left)** and β–β' (**c, right**) interfaces. Secondary structure elements participating in the interactions are indicated along the axes. The color key indicates the correspondence between color and net changes in contact numbers.

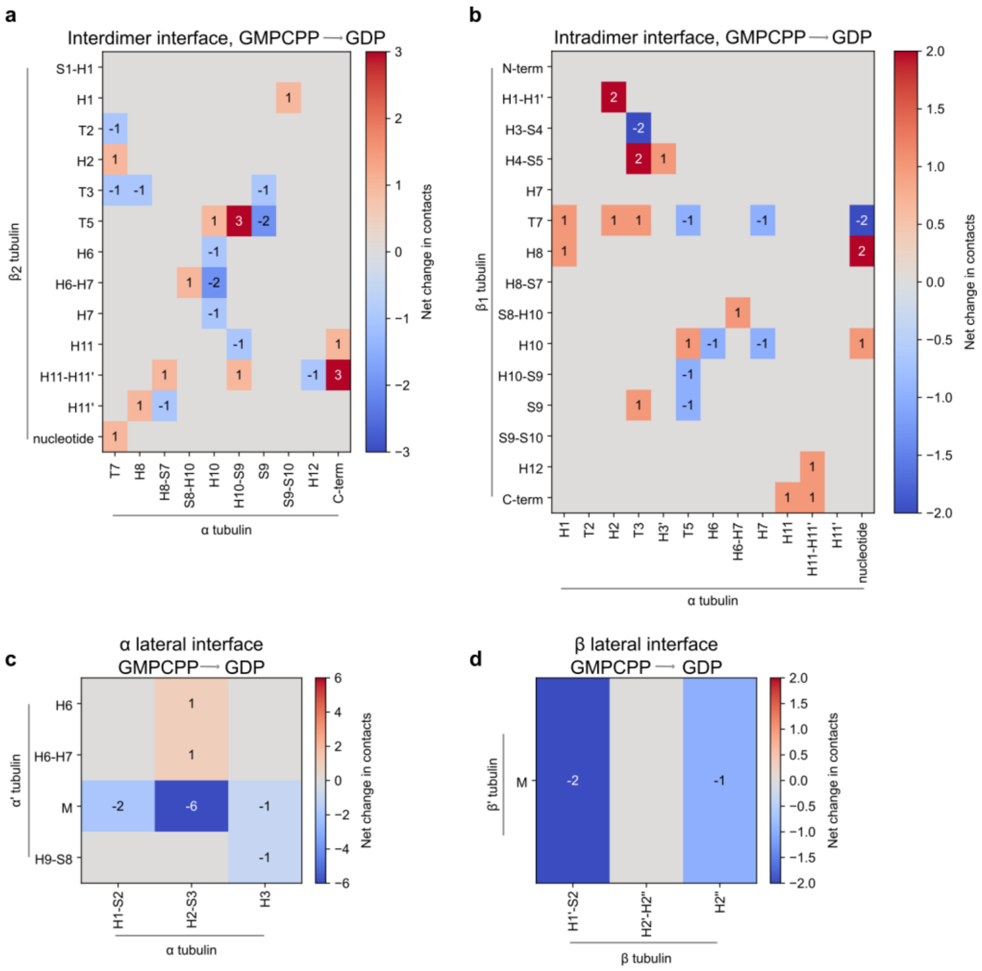


**Figure S8. Secondary structure contact balance for the GTP hydrolysis reaction.**

Heatmaps describing local changes in contact numbers at the longitudinal interdimer (**a**), longitudinal intradimer (**b**), lateral α–α' (**c**), and lateral β–β' (**d**) interfaces. Secondary structure elements participating in the interactions are indicated along the axes. The color key indicates the correspondence between color and net changes in contact numbers.

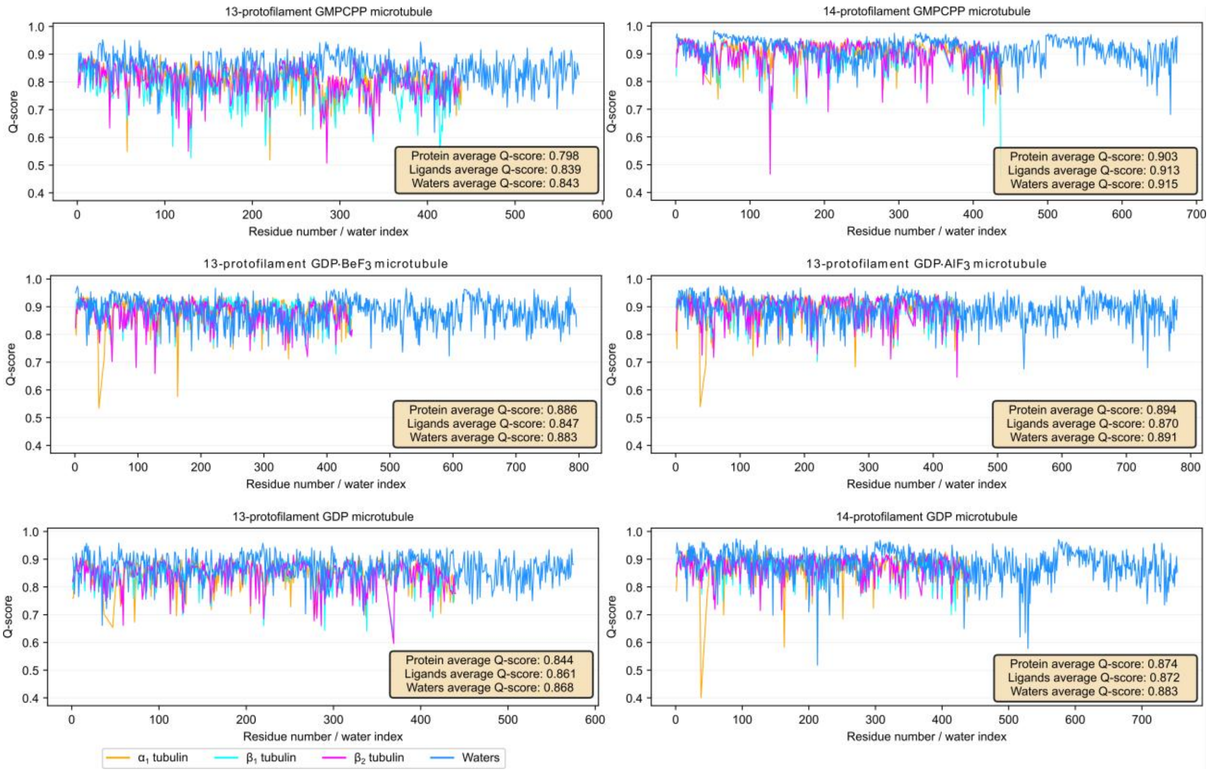


**Figure S9. Q-score analysis of microtubule structures.** Plots show Q-score values (y-axis) for residues in the $\alpha_1$- (orange), $\beta_1$- (light blue), and $\beta_2$-tubulin (purple) subunits, as well as for modeled water molecules associated with the three subunits (dark blue), plotted as a function of residue number or water index (x-axis). Q-score values for the nucleotides are shown in the inset at the bottom right of each subunit plot.

**Legends Supplementary Movies**

**Movie S1: Lattice and interface transitions.** First, lattice-scale transformations of 13-pf models are shown in cartoon representation, showcasing clear lattice compaction at the GMPCPP to GDP·$BeF_3^-$ transition, followed by very subtle lateral slide of the $\alpha_1$ tubulin on $\beta_2$ tubulin during the GDP·$BeF_3^-$ to GDP·$AlF_3$ transition, followed by an opposite movement during the GDP·$AlF_3$ to GDP transition. Next, sidechain movements during the reaction are shown, centered on interdimer and intradimer interface. Lastly, lateral interface transformations are shown, including modelled water molecules and hydrogen-bonding network. All transitions shown as morphs calculated by ChimeraX.

**Movie S2: GTP hydrolysis.** Detailed view of the catalytic site and its transformations throughout the GTP hydrolysis reaction. Protein residues in immediate vicinity of $\gamma$-phosphate and $Mg^{2+}$-coordinated waters shown. Water molecules within 6 Å of $Mg^{2+}$ shown as red spheres. Transitions between nucleotide states calculated as morphs in ChimeraX.
